# Persistence and Free Chlorine Disinfection of Human Coronaviruses and Their Surrogates in Water

**DOI:** 10.1101/2024.01.16.575911

**Authors:** Mengyang Zhang, Michelle Wei Leong, William A. Mitch, Catherine A. Blish, Alexandria Boehm

**Affiliations:** Department of Civil and Environmental Engineering, School of Engineering and Doerr School of Sustainability, Stanford University, Stanford, CA 94305, United States; Department of Medicine, Stanford University School of Medicine, Stanford, CA 94305, United States

**Keywords:** Human coronavirus, persistence, disinfection, drinking water

## Abstract

The COVID-19 pandemic illustrates the importance of understanding the behavior and control of human pathogenic viruses in the environment. Exposure via water (drinking, bathing, and recreation) is a known route of transmission of viruses to humans, but the literature is relatively void of studies on the persistence of many viruses, especially coronaviruses, in water and their susceptibility to chlorine disinfection. To fill that knowledge gap, we evaluated the persistence and free chlorine disinfection of human coronavirus OC43 (HCoV-OC43) and its surrogates, murine hepatitis virus (MHV) and porcine transmissible gastroenteritis virus (TGEV), in drinking water and laboratory buffer using cell culture methods. The decay rate constants of human coronavirus and its surrogates in water varied depending on virus and water matrix. In drinking water prior to disinfectant addition, MHV showed the largest decay rate constant (2.25 day^-1^) followed by HCoV-OC43 (0.99 day^-1^) and TGEV (0.65 day^-1^); while in phosphate buffer, HCoV-OC43 (0.51 day^-1^) had a larger decay rate constant than MHV (0.28 day^-1^) and TGEV (0.24 day^-1^). Upon free chlorine disinfection, the inactivation rates of coronaviruses were independent of free chlorine concentration and not affected by water matrix, though they still varied between viruses. TGEV showed the highest susceptibility to free chlorine disinfection with the inactivation rate constant of 113.50 mg^-1^ min^-1^ L, followed by MHV (81.33 mg^-1^ min^-1^ L) and HCoV-OC43 (59.42 mg^-1^ min^-1^ L).

**Importance:** This study addresses an important knowledge gap on enveloped virus persistence and disinfection in water. Results have immediate practical applications for shaping evidence-based water policies, particularly in the development of disinfection strategies for pathogenic virus control.

## Introduction

Individuals can acquire viral infections via exposure to contaminated water, through drinking, bathing, and recreating. Globally, at least 2 billion people use a drinking water source contaminated with feces; exposure to resultant microbial contamination can cause diarrhoeal diseases, acute respiratory diseases and various neglected tropical diseases.^1^ Waterborne viral pathogens are classified with moderate to high health significance by the World Health Organization (WHO) because they can have prolonged persistence in water supplies, moderate resistance to chlorine disinfection, and high relative infectivity.^2^

Human coronaviruses, which are single-stranded positive-sense RNA viruses from the family of *Coronaviridae*, have been circulating in human populations for decades. The first human coronavirus was isolated in 1965, while three highly pathogenic human coronaviruses emerged in the last 20 years: severe acute respiratory syndrome coronavirus (SARS-CoV-1), Middle East respiratory syndrome coronavirus (MERS-CoV), and SARS-CoV-2.^3,4^ The other four human coronaviruses (HCoVs) HCoV-229E, HCoV-OC43, HCoV-NL63, and HCoV-HKU1 are seasonal or endemic human coronaviruses, typically result in mild to moderate upper-respiratory tract illness, are responsible for 15-30% of common colds in adults, and represent the second most prevalent cause of the common cold.^5–7^ Notably, though HCoVs typically induce mild and self-limiting symptoms in immunocompetent adults, they could provoke lower respiratory tract inflammation and result in fatal outcomes among vulnerable populations such as infants, the elderly, pregnant women, and other immunocompromised individuals.^7–9^ HCoV-OC43 has also shown neuroinvasive potential, supporting observations of persistent replication in primary neuronal cell cultures and confirmed HCoV-OC43 neuroinvasive disease cases in humans.^10–13^ Regarding the global epidemiology of HCoVs, they exhibit seasonal patterns with peak activity typically occurring between December and March (Northern hemisphere) annually.^6,14^ The predominant HCoV species can vary by region, year, or age group, with HCoV-OC43 being the most commonly detected and peaking annually.^6,14,15^

During the COVID-19 pandemic, a number of studies were completed that investigated the environmental occurrence, persistence, and disinfection of HCoVs, including SARS-CoV-2, but these studies primarily focused on surfaces of fomite and food or in the air/aerosol droplets,^16–18^ as well as wastewater surveillance^19–21^. HCoVs have demonstrated the ability to survive in aerosols for hours to days, with persistence influenced by factors such as temperature and humidity.^16,18^ HCoVs also exhibit high persistence on fomites, with examples like HCoV-229E and SARS-CoV-2 surviving for 3-5 days on stainless steel surfaces.^16^ However, there are very limited studies on the persistence of HCoVs in water, particularly drinking water, and their susceptibility to chlorine disinfection.^22–24^

A few studies imply HCoVs could be persistent in water but susceptible to free chlorine disinfection in water but detailed evaluations are lacking. HCoV-229E could survive in dechlorinated tap water for ∼ 8 days (time to 99% inactivation, T99, at 23°C) and decayed significantly slower at a lower temperature (i.e., T_99_ >100 days at 4°C).^25^ Bivins et al.^26^ reported a T_99_ of ∼ 3.9 days for SARS-CoV-2 at 20°C in tap water had not been dechlorinated. Several other studies on the persistence of HCoVs and SARS-CoV-2 in water matrices were conducted but targeting the viral RNA which does not reflect the virus infectivity and usually shows higher persistence compared with intact, infectious viruses.^22,24^ As a common surrogate for enveloped virus, *Pseudomonas phage* Phi6 showed great free chlorine susceptibility at 23°C in phosphate-buffered saline (PBS) at pH 7.4 with an inactivation rate constant of 276.0 ± 30 mg^-1^ min^-1^ L.^27^ However, there has been no human coronavirus inactivation rate constant with exposure to free chlorine reported so far. Thus, there is a significant research gap regarding HCoVs persistence in drinking water and their susceptibility to free chlorine disinfection.

This study fills an important research gap on human coronavirus survival in water and their potential for chlorine disinfection. In particular, we demonstrate the persistence and free chlorine disinfection of infectious human coronaviruses and their surrogates in drinking water and laboratory buffer. HCoV-OC43 was selected as a representative human coronavirus because of its high prevalence in humans.^6,14,15^ We also selected murine hepatitis virus (MHV) and transmissible gastroenteritis virus (TGEV), which are mammalian coronaviruses infecting mice and pigs, respectively, and are common alternative viruses used in HCoV-related research.^28,29^ The decay rate constants of coronaviruses in water and their free chlorine disinfection kinetics were measured through cell culture-based assays. The results of this study will be of immediate use to inform water policies for pandemic preparedness and also highlight the necessity of selecting appropriate virus surrogates and water matrices in research on human pathogenic virus persistence and disinfection in water.

## Experimental Methods

### Overview

The persistence and free chlorine disinfection of human coronaviruses and their surrogates in drinking water were investigated by measuring their decay rate constants and free chlorine disinfection kinetics in drinking water. Factors including different viruses (HCoV-OC43, MHV, TGEV), water matrix (drinking water, phosphate buffer), and free chlorine concentration (mean value ± standard deviation: 1.31±0.06 mg L^-1^, 2.53±0.08 mg L^-1^) were investigated and statistically analyzed to identify the significant predictors for coronavirus decay and free chlorine disinfection kinetics in water. The concentration of infectious coronaviruses was quantified using a cell culture-based endpoint dilution assay known as the TCID_50_ method.

### Cells and virus propagation

Delayed brain tumor (DBT) cells were kindly provided by Dr. Krista Wigginton (University of Michigan, Ann Arbor, MI); swine testicular cells (ST) (CRL-1746, ATCC) and Vero E6 cells (CRL-1586, ATCC) were purchased from ATCC. All cell lines were maintained in Dulbecco’s modified Eagle’s medium (DMEM) supplemented with 10% fetal bovine serum (FBS) and 1% Penicillin-Streptomycin (P/S) and incubated at 37℃/5% CO_2_ until confluence. HCoV-OC43 and MHV were kindly provided by Dr. Jeffrey S. Glenn (Stanford University, Stanford, CA) and Dr. Krista Wigginton, respectively. For virus propagation, HCoV-OC43, MHV, and TGEV (VR-1995, ATCC) were inoculated to confluent Vero E6, DBT, and ST cells respectively, which were all maintained in serum free DMEM, at a multiplicity of infection (MOI) of 0.001-0.01. Infected cells were incubated at 37℃/5% CO_2_ until significant cytopathic effect (CPE) was observed, which usually took 6, 1, and 2 days for HCoV-OC43, MHV, and TGEV respectively. Virus suspension was collected and stored at -20 °C until further virus concentration and purification (see **Virus purification**).

### Virus purification

The presence of cell culture components and culture medium in virus stocks during the virus propagation process could cause significant bias in virus inactivation studies, by, for example, introducing artificial chlorine demand to experimental water matrices in virus seeding steps.^30^ In particular, DMEM is enriched with amino acids and vitamins (**Table S1**), which react with free chlorine rapidly^30^. To minimize the introduction of constituents with significant chlorine demand, all coronavirus stocks were purified through an ultracentrifugation-Amicon filtration process. First, crude coronavirus suspension was centrifuged at 4,000 ×g for 10 minutes. Then, the supernatant was filtered through 0.22 µm-pore size membranes to remove cell debris, and 120 mL of the filtrate was processed by ultracentrifugation at 24,000 rpm for 2.5 hours with a 30% of sucrose cushion. The coronavirus pellets were rinsed with phosphate buffer (10 µM, pH = 7.1) once and then resuspended in 1 mL autoclaved phosphate buffer. The 1 mL coronavirus suspension was then washed twice using AmiconUltra (Merck Millipore) with 10 mL phosphate buffer. The result was approximately 150-200 µL of purified coronavirus stock. The purified coronavirus stocks were stored at -80 °C for up to two weeks until used in persistence and free chlorine disinfection experiments.

### Persistence experiments

We obtained water samples from the Vineyard Water Treatment Plant in Sacramento, California, USA, which produces potable water for the surrounding communities. Water samples were collected right after the sedimentation process and before chlorine addition. Water quality parameters are provided in **Table S2**. The water samples were filtered through 0.7 µm pore size glass fiber membranes (Whatman) to mimic the filtration process in the Vineyard Water Treatment Plant and then stored at -80 °C for up to a year until use. This water is subsequently referred to as “drinking water”.

290 µL of purified HCoV-OC43, MHV, and TGEV stocks were spiked into either 2,610 µL of drinking water or autoclaved phosphate buffer at a ratio of 1:10 separately to achieve final coronavirus concentrations of ∼ 10^5^-10^6^ TCID_50_ mL^-1^; incubated at room temperature (22°C) for up to 30 days. During the incubation period, reactors were kept on a sample mixer (HulaMixer^TM^) and rotated vertically at 1 rpm. 60 µL of subsamples were collected on days 0, 1, 2, 3, 5, 7, 11, 17, and 30, and tested immediately for the TCID_50_ to quantify the concentration of infectious coronaviruses (see below **TCID_50_ for virus titration**). Two reactors were used for each coronavirus (experimental duplicates) and two subsamples were collected at each time point from each reactor (technical duplicates), resulting in four replicates for each time point.

The decay rate constants (*k_decay_*) were estimated by fitting a first-order decay model to the corresponding persistence data from four replicates, while excluding the data under the lower limit of detection (LoD):

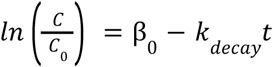

where *C* is the average infectious concentration of each coronavirus at each time point in each water matrix (TCID_50_ mL^-1^), *C_0_* is the average infectious concentration of each coronavirus at time 0 in each water matrix (TCID_50_ mL^-1^), β is the intercept, *k_decay_* is the first-order decay rate constant for each condition (day^-1^), and *t* is the incubation time (day). *k_decay_* and its standard error were determined using the function “lm” in R (version 4.2.2) and Rstudio (Version 2022.12.0+353), and the goodness of fit of the first-order decay model was assessed through examination of the coefficient of determination (R^2^).

### Free chlorine disinfection

Previous work has indicated that enveloped viruses are particularly susceptible to free chlorine exposure^27^, and our own pilot experiments suggested that coronaviruses are as well (data not shown). We therefore created a modified lab-scale continuous quench-flow system (**Figure S1**) to investigate the free chlorine disinfection of coronaviruses. This system allows one to measure reaction kinetics for reactions that are too fast to observe in batch reactors.^31^ In brief, free chlorine working solution and coronavirus working solution were pumped through a six-channel syringe pump (Masterflex^®^), for which the flow rate was set as 30 mL h^-1^, and then mixed in a PEEK micro static mixing tee (IDEX Health & Science). The reacting mixture flowed through a sample loop of pre-defined length and then was quenched by a four-fold molar excess of ascorbic acid (A61-100, Fisher Scientific) relative to the initial free chlorine. The ascorbic acid was dissolved directly in DMEM to minimize the requirement of dilution of coronaviruses at the end of the disinfection experiments. Reaction times were controlled by using different sample loop lengths. As the flow rate of reacting mixture flowing through the sample loop is fixed (i.e., 60 mL h^-1^), the contact time of the free chlorine disinfection process is only dependent on the length of the sample loop, and can be calculated by:

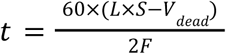

where *t* is the contact time (min), 60 represents 60 min h^-1^, *L* is the length of the sample loop (cm), *S* is the surface area of the sample loop (1/16” OD ⋅ 0.010” ID tubing, 0.0005 cm^2^), *V_dead_* is the dead volume in the micro static mixing tee (0.00095 cm^3^), and *F* is the flow rate of the syringe pump (30 mL h^-1^).

The free chlorine working solution was prepared by diluting a NaOCl stock solution (Sigma-Aldrich) in 10 mM autoclaved phosphate buffer at pH of 7.1, where HOCl and OCl^-^ are at a ratio of 72:28 in molar concentrations. Phosphate buffer was selected instead of phosphate buffer saline since high levels of chloride can affect speciation of free chlorine and thus the reaction kinetics of some organic compounds.^32^ The HCoV-OC43, MHV, and TGEV working solutions were prepared by diluting purified HCoV-OC43, MHV, and TGEV stocks at least 1:100 either in drinking water or phosphate buffer to minimize chlorine demand introduced by DMEM residual. The pH of the mixture of drinking water and free chlorine working solution at 1:1 ratio was 7.1. Therefore, the pH was consistent between the chlorine disinfection experiments conducted using drinking water or phosphate buffer. Two free chlorine concentrations of approximately 1.30 mg L^-1^ (referred to as “low concentration” subsequently) and 2.50 mg L^-1^ (referred to as “high” concentration” subsequently) were used to investigate the impact of free chlorine concentration on coronavirus chlorine disinfection kinetics in water. The concentration of free chlorine working solution was prepared as 2.60 mg L^-1^ and 5.00 mg L^-1^ as Cl_2_ to obtain the targeted final free chlorine concentration of 1.30 mg L^-1^ and 2.50 mg L^-1^ as Cl_2_ in the reacting mixture after mixing with the coronavirus working solution. The free chlorine concentration was measured by a modified *N*,*N*-diethyl-*p*-phenylenediamine colorimetric method (see **Supplemental Material** for details).^33^ The chlorine demand of coronavirus working solutions and the actual free chlorine concentration in the reacting mixture were measured before every free chlorine disinfection experiment. The chlorine demand of drinking water was negligible after exposure to the highest chlorine concentration and contact time (**Table S3**). The samples after free chlorine disinfection experiments were immediately tested using the TCID_50_ assay for the measurement of infectious virus concentration (see below for **TCID_50_ for virus titration**). For time 0 samples, coronavirus working solutions were mixed with pre-quenched free chlorine working solutions. Three independent experiments were conducted at each time point across all conditions shown in **Table S4**.

The inactivation rate constants (*k_inactivate_*) were estimated by fitting a pseudo-first-order model to the corresponding free chlorine disinfection data from 3 replicates while excluding the data under the lower limit of detection (LoD), since the chlorine concentration was constant:

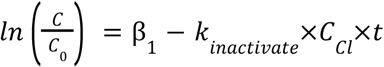

where *C* is the average infectious concentration of each coronavirus at each time point (TCID_50_ mL^-1^), *C_0_* is the average infectious concentration of each coronavirus at time 0 (TCID_50_ mL^-1^), β is the intercept, *k_inactivate_* is the pseudo-first-order inactivation rate contestant for each condition (mg^-1^ min^-1^ L), *C_Cl_* is the measured initial free chlorine concentration in each experiment (mg L^-1^, **Table S4**), and *t* is the contact time (min, **Table S4**). *k_inactivation_* and its standard error were determined using the function “lm” in Rstudio for data of *ln*(*C/C_0_*) versus CT (*C_cl_*×*t*), and the fitness of the pseudo-first-order decay model was assessed by determining the coefficient of determination (R^2^).

### TCID_50_ for virus titration

A endpoint dilution assay of TCID_50_ was used to measure coronavirus infectivity.^34^ In brief, HCoV-OC43, MHV, and TGEV samples were serially diluted 10-fold (i.e., 10^0^-10^-7^) with DMEM supplemented with 1% P/S and 100 µL of the diluted samples was inoculated to confluent Vero E6, DBT, and ST cells respectively, which were all maintained in 96-well plates and rinsed with serum free DMEM once. Five replicate wells were applied for each dilution. After incubation at 37°C/5% CO_2_ for 7, 2, or 4 days for HCoV-OC43, MHV, and TGEV, respectively, when significant cytopathic effect was observed, supernatant was removed and the cells were stained with 1% crystal violet (HT901, Sigma-Aldrich, diluted with ethanol) for 10 min^35^. After the crystal violet was rinsed away using tap water, the number of positive wells that were transparent or had large clearings in the cell lawn were counted and the infectious concentration of coronaviruses was calculated as TCID_50_ mL^-1^ as per Spearman–Kärber method^36^. The LoD was defined as the concentration where only one out of five wells was scored as positive at the lowest dilution, corresponding to 5.0 TCID_50_ mL^-1^ in this study.

### Statistical Analysis

Analysis of covariance (ANCOVA) was applied to test whether two decay or inactivation constants were significantly different using GraphPad Prism 9 software. For the persistence results, a total of 9 pairs of comparison was conducted via ANCOVA (HCoV-OC43/MHV, HCoV-OC43/TGEV, and MHV/TGEV in drinking water or phosphate buffer; drinking water/phosphate buffer condition for each virus); for the disinfection results, a total of 21 pairs of comparison was conducted via ANCOVA (drinking water/phosphate buffer condition with low/high concentration of free chlorine for each virus; HCoV-OC43/MHV, HCoV-OC43/TGEV, and MHV/TGEV for merged condition). The null hypotheses of no difference between two decay or inactivation rate constants were rejected if p values were <0.05.

All data from this study (time series of measured viral concentrations, calculated *k_decay_* and *k_inactivation_*values, and statistical analysis results) are available in the Stanford Digital Repository DOI: https://purl.stanford.edu/tm847rm8309.

## Results

### Persistence of coronaviruses in water

Purified HCoV-OC43, MHV, and TGEV stocks were spiked into either drinking water prior to disinfectant addition or phosphate buffer and incubated for 30 days at room temperature to observe their decay rates. As shown in **Figure 1** and **Table S5**, the times required for a 99% reduction in infectivity (T_99_) of MHV and TGEV in drinking water were notably shorter, with MHV taking 2.0 days and TGEV taking 7.1 days, whereas a significantly longer duration of 16.5 days for MHV and 19.4 days for TGEV was necessary for achieving the same level of reduction when they were in phosphate buffer. For HCoV-OC43, the T_99_ values were 4.6 days in drinking water and 9.0 days in phosphate buffer.

**Figure 1.**
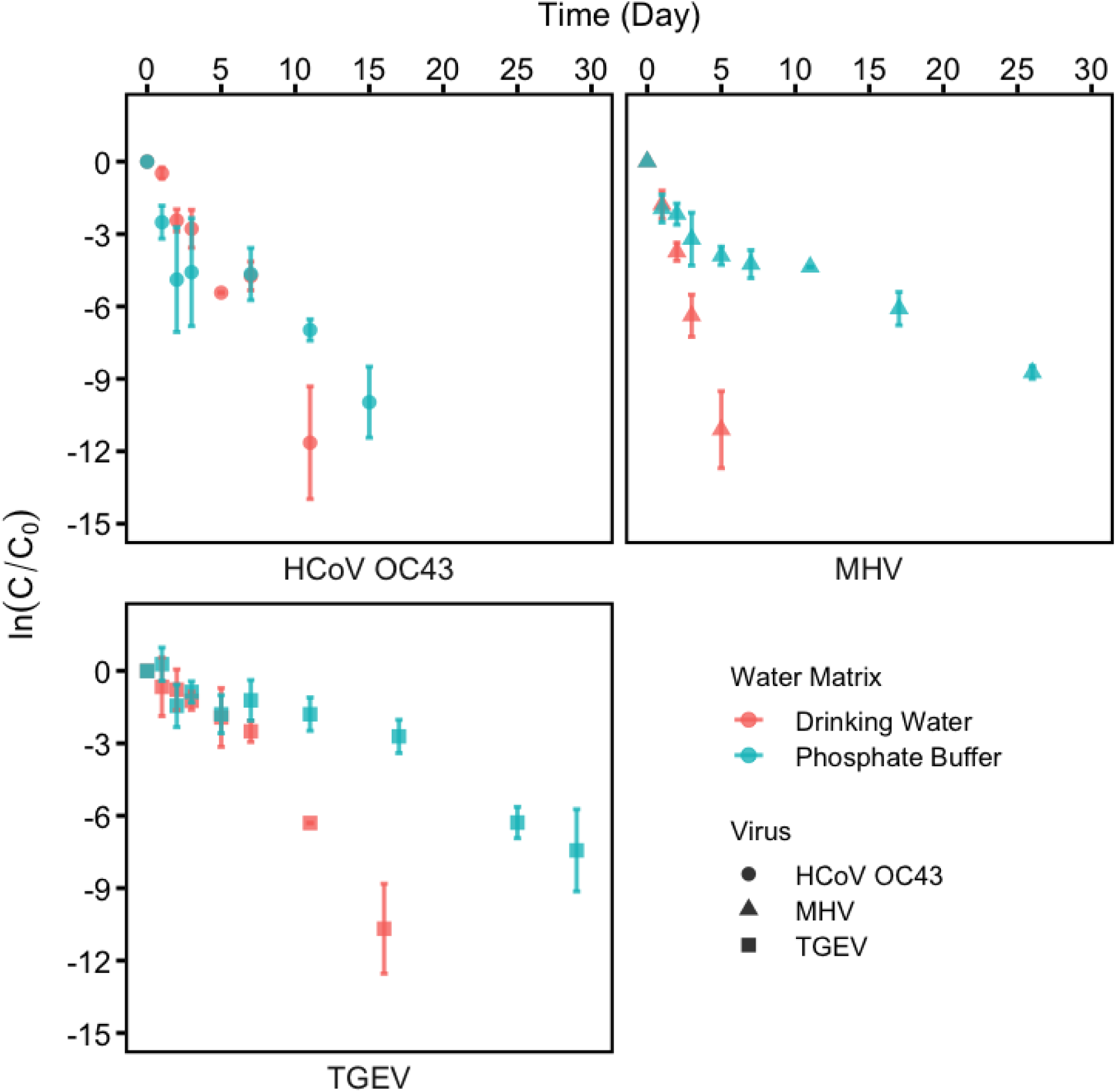
Decay of coronaviruses in drinking water and phosphate buffer. Error bar represents standard deviation. n=4 for each data point.

The decay rate constants *k_decay_* were estimated by fitting a first-order model (**Figure 2)**. HCoV-OC43, MHV, and TGEV all exhibited significantly higher decay rate constants in drinking water than in phosphate buffer, i.e., around 1.9-8.1 fold higher (p<0.05, **Tables S5** and **S6**). In drinking water, MHV demonstrated the highest decay rate constant with a *k_decay_* of 2.25±0.09 day^-1^, while in phosphate buffer, HCoV-OC43 had the largest decay rate constant of *k_decay_* of 0.51±0.10 day^-1^ (**Table S5**). Notably, TGEV has the smallest decay rate constants in both drinking water and phosphate buffer, i.e., 0.65±0.06 day^-1^ and 0.24±0.02 day^-1^, respectively. Decay rate constants for MHV and TGEV in phosphate buffer were not significantly different (0.28±0.03 day^-1^ and 0.24±0.02 day^-1^ respectively, p=0.5724).

**Figure 2.**
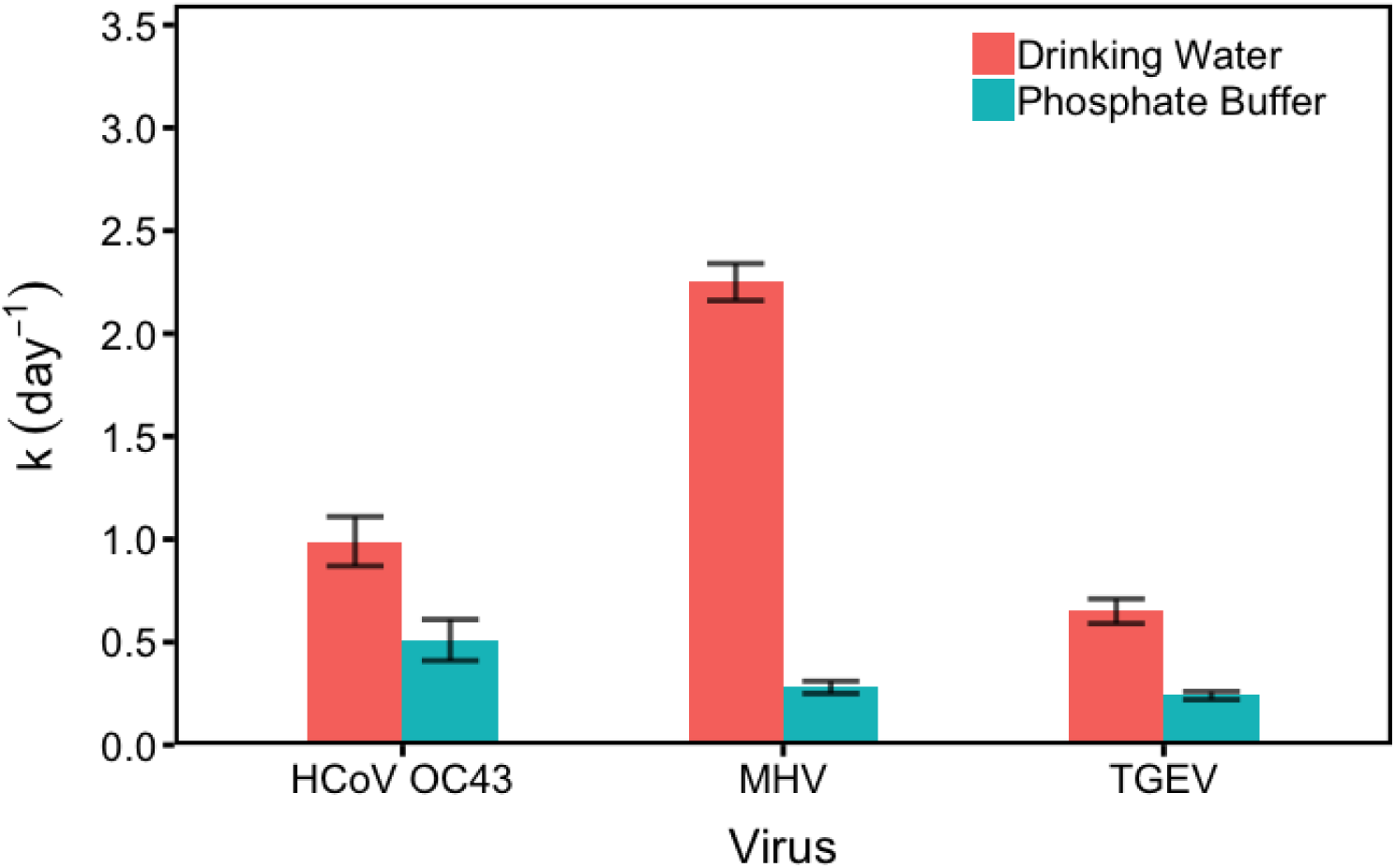
First-order decay rate constants of coronaviruses in drinking water and phosphate buffer. Error bar represents standard error.

### Free chlorine disinfection kinetics of coronaviruses

The susceptibility of coronaviruses to disinfection by free chlorine was evaluated with initial free chlorine concentrations of 1.31±0.06 and 2.53±0.08 mg L^-1^, which fall in the range of 0.5-4 mg L^-1^ recommended by EPA for both effective disinfection and safety in drinking water.^2^ As shown in **Figure 3**, HCoV-OC43 was inactivated by ∼ 2.1 log_10_ after exposure to free chlorine with a CT value of 0.08 mg min L^-1^ (i.e., free chlorine concentration × contact time: 1.30 mg L^-1^ of free chlorine × 3.5 s or 2.50 mg L^-1^ of free chlorine × 1.8 s) in drinking water and phosphate buffer. MHV and TGEV demonstrated greater inactivation compared to HCoV-OC43 when exposed to an equivalent CT value of 0.08 mg min L^-1^, exhibiting reductions in virus titer by ∼ 2.8 log_10_ and 4.0 log_10_, respectively.

**Figure 3.**
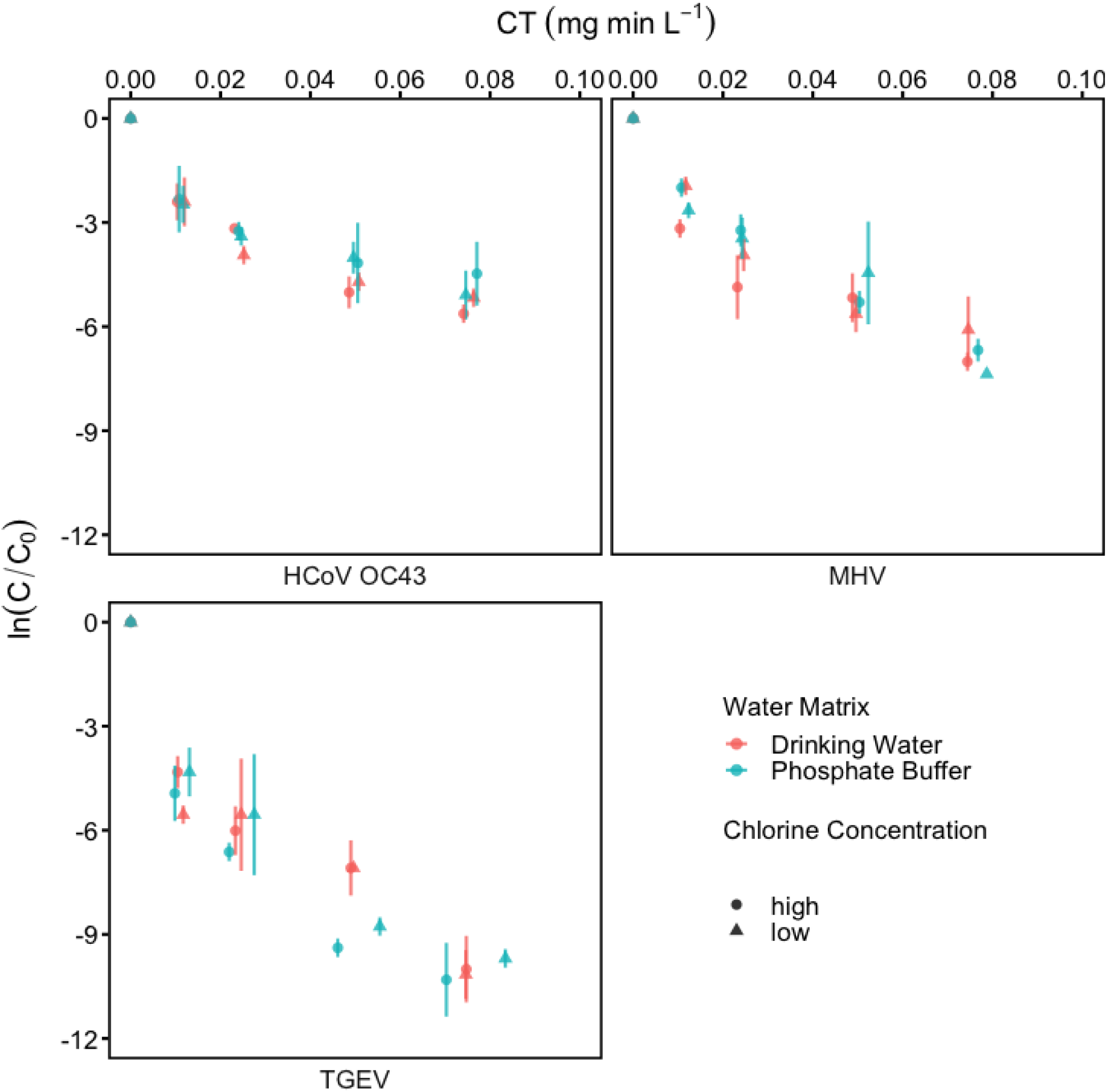
Inactivation of coronaviruses by free chlorine in drinking water and phosphate buffer. Water Matrix represents the virus working solution matrices in free chlorine disinfection experiments; High (2.53±0.08 mg L^-1^) or low (1.31±0.06 mg L^-1^) Chlorine Concentration represents the initial free chlorine concentration used in free chlorine disinfection experiments. Error bar represents standard deviation. n=3 for each data point.

Although inactivation was faster at higher free chlorine concentrations, second order inactivation rate constants (*k_inactivate_*; mg^-1^ min^-1^ L estimated from the pseudo-first-order model were not significantly different either with different initial free chlorine concentrations or for different water matrices (p>0.05, **Figure 4**, **Table S7**). Thus, the inactivation rate constants for each virus were estimated through the consolidation of data obtained from all disinfection experiments for each virus, including exposure to low and high concentrations of free chlorine in both drinking water and phosphate buffer, followed by fitting the data to a pseudo-first-order model (**Table S8**). The inactivation rates of coronaviruses by free chlorine varied significantly between viruses (p<0.05, **Table S9**). TGEV had the highest inactivation rate constant of 113.50 mg^-1^ min^-1^ L followed by MHV (81.33 mg^-1^ min^-1^ L) and HCoV-OC43 (59.42 mg^-1^ min^-1^ L).

**Figure 4.**
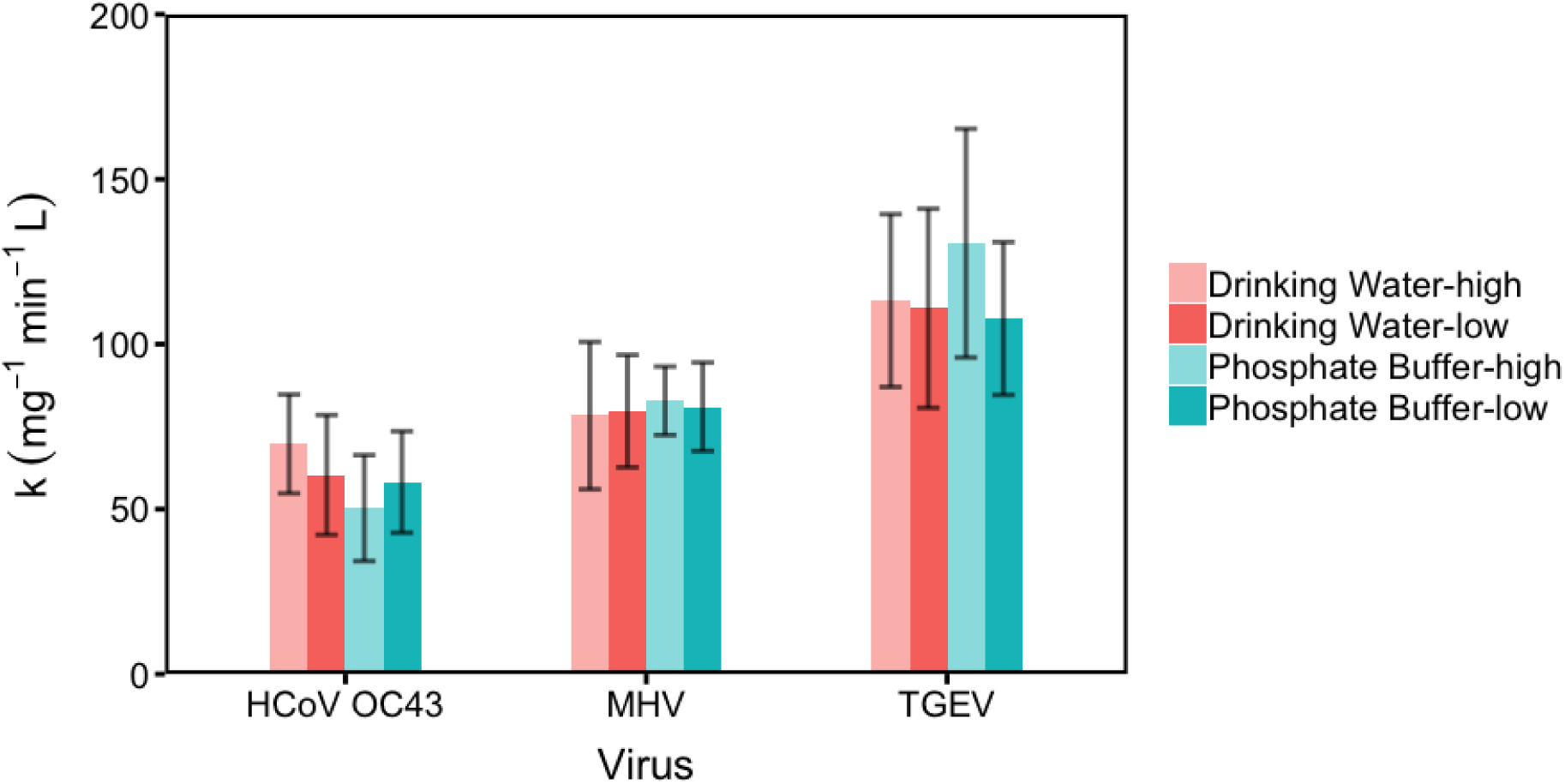
Pseudo-first-order inactivation rate constants of coronaviruses by free chlorine disinfection. Drinking Water or Phosphate Buffer represents the virus working solution matrices in free chlorine disinfection experiments; High and low represents the initial free chlorine concentration of 2.53±0.08 mg L^-1^ and 1.31±0.06 mg L^-1^ used in free chlorine disinfection experiments, representatively. Error bar represents standard error.

## Discussion

This study demonstrated that HCoV-OC43 and surrogate viruses MHV and TGEV are persistent in water, and that the decay varied between water matrices and viruses. Prolonged survival of MHV, TGEV, and SARS-CoV in laboratory buffers or water has been observed in previous studies,^37,38^ and the decay rate constants reported here are within the range of those previously reported for viruses in water^23^. Previous work on virus persistence in water suggests that enzymatic activity, predation, particles, and the presence of solvents, detergents, and organic matter can accelerate virus inactivation;^25,37,39^ these were all absent or present in low concentrations in the simplified waters studied herein. The different decay rate constants of HCoV, MHV, and TGEV in water suggests that intrinsic properties of the viruses affect their stability in water. A meta-analysis of decay rate constants of enveloped viruses in water found that virus type significantly affected viral persistence.^40^ For example, they reported that Influenza A subtype H11N6 had larger decay rate constants than other influenza A subtypes; enveloped viruses from the *Togaviridae* family decayed slower than influenza A viruses and some viruses from the Rhabdoviridae and *Coronaviridae* families.^40^ As more data are collected on virus persistence in water, it is our hope that greater insights will be developed on how virus characteristics affect their decay.

HCoV-OC43, MHV, and TGEV all showed susceptibility to free chlorine disinfection (Figure 3). We tested both biphasic and pseudo-first-order models for the inactivation, and the pseudo-first-order model fits better than the biphasic model (see **Supplemental Material** for model equation) as illustrated by lower Akaike Information Criteria (AIC) values for all free chlorine disinfection conditions (**Table S10**). The pseudo-first-order inactivation rate constants (using the product of free chlorine concentration and contact time (CT) as the variable) of coronaviruses were independent of free chlorine concentration and not affected by water matrix, though they still varied between viruses. Lipids containing saturated fatty acids are widely recognized for their high resistance to oxidation by free chlorine,^41^ particularly in contrast to proteins and genomes.^27^ However, other research has demonstrated that lipids containing unsaturated fatty acids react rapidly with free chlorine.^42^ Previous work suggests that genome damage might be a major mechanism for free chlorine inactivation of viruses like hepatitis A virus and bacteriophage MS2,^43,44^ with adenosine and cytosine nucleobases being more chlorine-reactive^45^. However, the three viruses studied herein (HCoV-OC43, MHV, and TGEV) have similar proportions of chlorine-reactive bases in their genomes (71-74% as total chlorine-reactive bases (C+A+G), **Table S11**), which indicates that their differential susceptibility to free chlorine might not solely be driven by their genome composition. Chlorine-induced protein damage is another significant contributor to virus inactivation by free chlorine.^27,43^ Specifically, the inactivation of *Pseudomonas phage* Phi6, a common surrogate for enveloped viruses, by free chlorine was reportedly driven by the oxidation of nucleocapsid proteins and proteins in the polymerase complex; free chlorine was shown to readily penetrate the envelope membrane.^27^ Methionine (Met) and cysteine (Cys) are two of the most chlorine-reactive amino acid residues in viral proteins, with second-order reaction rate constants of 3.8×10^7^ M^-1^ s^-1^ and 3.0×10^7^ M^-1^ s^-1^, respectively.^46^ Notably, Phi6 membrane proteins predominantly lack these amino acids. In contrast, the presence of Met and Cys is pervasive among proteins in HCoV-OC43, MHV, and TGEV (**Table S12**). In the present study, the relationship between the absolute counts of Met and Cys in viral proteins and virus inactivation rate constants by free chlorine varied depending on the protein’s location. For example, a negative correlation (correlation coefficient r=-0.74, p<0.001) was observed between Met counts in spike proteins and virus inactivation rate constants, while a positive correlation (r=0.83, p<0.001) was found between Cys counts in membrane proteins and virus inactivation rate constants (**Table S13**). The varied correlations between Met and Cys counts in viral proteins and virus inactivation rate constants by free chlorine suggests the protein composition might not be the only factor driving the protein degradation by free chlorine. Both Ye et al^27^ and Sigstam et al^43^ emphasized the significant role of solvent-accessible surface area of individual amino acids in the degradation of proteins or peptides, i.e., slower degradation of Met- and Cys-containing peptides was observed when they were surrounded by other amino acid residues. Remarkably, despite the striking structural similarities among HCoV-OC43, MHV, and TGEV, it is noteworthy that only HCoV-OC43 exhibits the expression of the Hemagglutinin-esterase protein HE,^47^ which is located on the enveloped membrane and might potentially play a role in the inactivation of HCoV-OC43 due to the presence of Met and Cys. Nevertheless, the oxidation kinetics of individual proteins or peptides in HCoV-OC43, MHV, and TGEV remain unclear, calling for further investigation.

It is increasingly recognized that various viruses, even those closely related or possessing similar structural features, can display markedly divergent responses when exposed to environmental water or disinfectants.^43,48^ However, the underlying mechanisms driving these distinct viral responses remain elusive, necessitating further research efforts. For example, comprehensive analyses such as proteomic and lipidomic assessments, as well as investigations into the degradation kinetics of individual virus peptides, could be systematically conducted on viruses after exposure to environmental stressors or disinfectants. Moreover, advanced genetic approaches, like sequencing, may provide insight into resultant genome damage, i.e., by identifying and quantifying damage in specific regions of the genome.

Experimental artifacts can also significantly impact results and potentially lead to misinterpretations in studies of virus disinfection. For instance, the impurities present in virus stocks used during experiments can introduce additional oxidant demand, complicating virus disinfection investigations.^30^ The presence of chloride in commonly used laboratory buffers, such as PBS buffer, may enhance the degradation of viral genomes when exposed to free chlorine, as it can form more effective halogenating agents than HOCl (e.g., Cl_2_).^49^ Moreover, given that the choice of host cell line is indispensable to virus culture and titration, the observed kinetics of enterovirus inactivation by free chlorine have been noted to be host cell-dependent.^50^ Discrepancies in inactivation kinetics of rotavirus during free chlorine disinfection were also observed in experiments using mice model versus cell culture model.^41^ These considerations underscore the importance of meticulous experimental design in future virus research to ensure accurate and meaningful results.

### Environmental Implication

This study addresses a significant knowledge gap by investigating the behavior of HCoVs in water, particularly focusing on their persistence and disinfection potential. While extensive research has been conducted on non-enveloped viruses in water,^51^ limited information exists regarding enveloped viruses like coronaviruses^23,40^. With the ongoing emergence of new enveloped viruses with pandemic potential, understanding how enveloped viruses like HCoVs behave in water can inform the development of water treatment and disinfection methods to mitigate the spread of infectious diseases, particularly when waterborne transmission is a concern. Advanced research and adaptive management strategies are critical to stay ahead of emerging threats.

Our research on the disinfection of HCoVs by free chlorine in water has immediate practical applications for shaping evidence-based water policies, particularly in the development of disinfection strategies for pathogenic virus control. With a CT value of 0.08 mg min L^-1^ of free chlorine, which can achieve at least a 2 log_10_ reduction in virus infectivity, our findings align with the concept of low resistance of pathogenic viruses to free chlorine (CT<1 mg min L^-1^ for 99% inactivation) as defined by WHO criteria^2^or typical free chlorine residual concentrations (∼ 0.5-1.5 mg/L), 2 log_10_ inactivation would be achieved in <20 seconds. Note that slower inactivation kinetics are expected for chloramines than for free chlorine disinfection. However, utilities employing chloramines for maintenance of a distribution system residual frequently incorporate free chlorine contact times prior to ammonia addition to form chloramines for distribution. Even where only short free chlorine contact times are employed (e.g., 3 min), our results suggest the potential for significant viral inactivation. While many utilities have switched from free chlorine. This information is pertinent not only to the safety of potable water but also for safeguarding recreational water sources like swimming pools, as both are integral to public health. The EPA recommends a range of 0.5-4 mg L^-1^ of free chlorine for both effective disinfection and safety in drinking water,^2^ while the CDC advocates for maintaining a minimum free chlorine concentration of 1 mg L^-1^ in swimming pools, and at least 3 mg L^-1^ in hot tubs and spas,^52^ underscoring the practical significance of our research in promoting public health and safety.

Furthermore, this research underscores the critical importance of meticulous selection of appropriate virus surrogates and environmentally relevant matrices, when assessing the persistence of human pathogenic viruses in the environment and their resistance to disinfectants. The significant variations in both persistence and resistance to free chlorine observed among HCoV-OC43, MHV, and TGEV herein highlight that even closely related or structurally similar viruses can exhibit markedly distinct behavior when exposed to environmental conditions and disinfectants. The conventional practice of using a universal surrogate for multiple viruses within the same family may no longer be applicable, a viewpoint supported by other researchers as well^43,48^. For instance, Aquino de Carvalho et al.^48^ advocate for the necessity of working with the virus of interest in environmental persistence studies, after their evaluation of Phi6 as a surrogate for enveloped viruses. Many previous studies on viruses in the environment have traditionally been carried out using laboratory buffers or artificial media. However, it is important to recognize that real environmental matrices are considerably more complex, encompassing not only a great variety of particles, solvents, detergents, and organic materials but also microbial communities, which all might impact the virus infectivity, persistence, and disinfection in the environment. This emphasizes the need for a more comprehensive approach to studying viruses in their true environmental contexts.

## Acknowledgment

This work was supported by the Stanford Woods Institute for the Environment.

