## Supporting material for "Persistence and Free Chlorine Disinfection of Human Coronaviruses and Their Surrogates in Water"

**Supplemental Material**

Mengyang Zhang,[1] Michelle Wei Leong,[2] William A. Mitch,[1] Catherine A. Blish,[2] Alexandria Boehm[1*]

[1] Department of Civil and Environmental Engineering, School of Engineering and Doerr School of Sustainability, Stanford University, Stanford, CA 94305, United States

[2] Department of Medicine, Stanford University School of Medicine, Stanford, CA 94305, United States

* Corresponding Author:

*Environmental Science and Technology*

13 Tables, 1 Figure, and 19 Pages

**Modified *N,N*-diethyl-*p*-phenylenediamine (DPD) colorimetric method for free chlorine quantification**

The DPD solution was prepared by dissolving 0.1 g of DPD in 5 mL of 0.1 N $H_2SO_4$ and stored at 4 °C. For quantifying free chlorine, the reaction was conducted by mixing 40 µL of the sample, 10 µL of 100 mM phosphate buffer (pH 6), and 30 µL of the DPD solution, and then transferred to a transparent 96-well plate as 80 µL mixed solution per well for each sample. Multiple reactions could be prepared simultaneously using a multi-channel pipettor, which also minimized the technical difference between samples. The absorbance was measured at 560 nm with a Microplate Reader (Promega, GloMax® Discover). The standard curve of free chlorine was obtained by quantifying the absorbance of serially diluted chlorine standard solutions (27.584 ± 0.18 mg $L^{-1}$ as free chlorine, HACH, 2630020) after reaction with DPD at each time of free chlorine disinfection. For the measurement of the chlorine demand of coronavirus working solutions or filter drinking water sample, a negative control of chlorine demand free phosphate buffer was also prepared simultaneously using a multi-channel pipettor to further adjust the standard curve and minimize the technical difference between different batches of tests.

**Biphasic model for free chlorine disinfection**

A biphasic model was applied for the fitting of free chlorine disinfection data following the equation:

$$ln\left(\frac{C}{C_0}\right) = A_1 * exp(-B_1 * CT) + A_2 * exp(-B_2 * CT)$$

where $C$ is the average infectious concentration of each coronavirus at each time point ($TCID_{50}$ $mL^{-1}$); $C_0$ is the average infectious concentration of each coronavirus at time 0 ($TCID_{50}$ $mL^{-1}$); $A_1$ and $A_2$ are the amplitudes for the first and second exponential terms, respectively; $B_1$ and $B_2$ are the decay constants for the first and second exponential terms, respectively; CT is the product of ($C_{Cl}$×$t$), while $C_{Cl}$ is the measured initial free chlorine concentration in each experiment (mg $L^{-1}$, **Table S4**), and $t$ is the contact time (min, **Table S4**). The function “nls” in Rstudio was used for the fitting of the biphasic model.

**Correlation testing**

A correlation matrix was computed to analyze the correlations between the absolute counts of chlorine-reactive amino acids in viral proteins and virus inactivation rate constants by free chlorine, by using the function of “chart.Correlation” in R (version 4.2.2) and Rstudio (Version 2022.12.0+353). In the plot, the distribution of each variable is shown on the diagonal; the bivariate scatter plots with a fitted line are displayed on the bottom of the diagonal; the value of the correlation plus the significance level as stars are displayed on the top of the diagonal. “***”, “**”, “*”, “.”, and “ ” represent p-values of 0, 0.001, 0.01, 0.05, 0.1, and 1 respectively. The absolute counts of chlorine-reactive amino acids, including Met (Methionine), Cys (Cysteine), His (Histidine), Trp (Tryptophan), Lys (Lysine), and Tyr (Tyrosine), were obtained by counting the number of "M", "C", "H", "W", "K", "Y" in viral genome sequences respectively in Rstudio. The inactivation rate constants (*k*) were estimated by fitting a pseudo-first-order model to the corresponding free chlorine disinfection data and determined using the function “lm” in Rstudio as described in the main manuscript. Each *k* was estimated from one distinct replicate dataset in free chlorine disinfection experiment conditions, including inactivation of HCoV-OC43, MHV or TGEV in drinking water or phosphate buffer by low or high concentration of free chlorine.

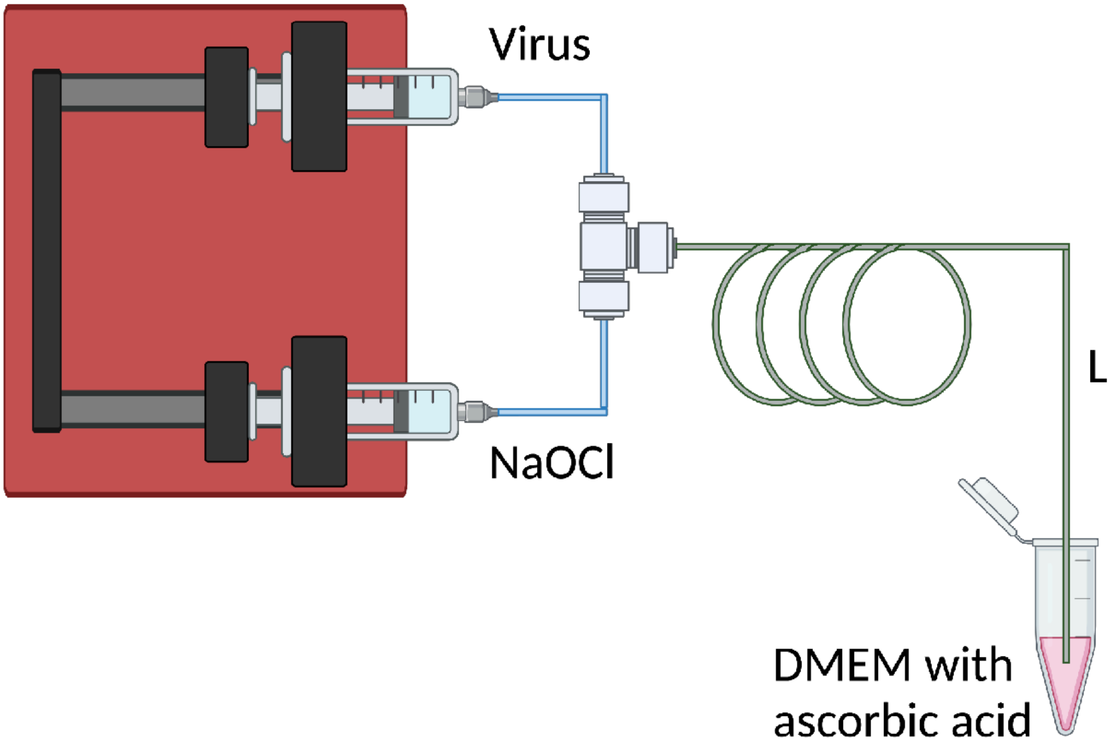


**Figure S1**. Lab-scale continuous quench-flow system for free chlorine disinfection experiment. L represents the length of the sample loop (cm).

**Table S1.** DMEM formula.

https://www.corning.com/catalog/cls/documents/formulations/CLS-CG-BR-001-DMEM-Formulations.pdf

| Units | mg $L^{-1}$ | | mg $L^{-1}$ | | mg $L^{-1}$ | | mg $L^{-1}$ |
|---|---|---|---|---|---|---|---|
| **Inorganic Salts** | | **Amino Acids** | | **Vitamins** | | **Other** | |
| $CaCl_2$ (anhydrous) | 200 | L-Arginine • HCl | 84 | D-Calcium pantothenate | 4.00 | D-Glucose | 4500.00 |
| $Fe(NO_3)_3 \cdot 9H_2O$ | 0.1 | L-Cystine • 2HCl | 62.57 | Choline chloride | 4.00 | Phenol red • Na | 15.00 |
| KCl | 400 | L-Glutamine | 584 | Folic acid | 4.00 | Sodium pyruvate | 110.00 |
| $MgSO_4$ (anhydrous) | 97.7 | Glycine | 30 | *i*-Inositol | 7.20 | | |
| NaCl | 6400 | L-Histidine •HCl • $H_2O$ | 42 | Nicotinamide | 4.00 | | |
| $NaH_2PO_4 \cdot H_2O$ | 125 | L-Isoleucine | 104.8 | Pyridoxine • HCl | 4.00 | | |
| $NaHCO_3$ | 3700 | L-Leucine | 104.8 | Riboflavin | 0.40 | | |
| | | L-Lysine • HCl | 146.2 | Thiamine • HCl | 4.00 | | |
| | | L-Methionine | 30 | | | | |
| | | L-Phenylalanine | 66 | | | | |
| | | L-Serine | 42 | | | | |
| | | L-Threonine | 95.2 | | | | |
| | | L-Tryptophan | 16 | | | | |
| | | L-Tyrosine • 2Na • $2H_2O$ | 103.79 | | | | |
| | | L-Valine | 94 | | | | |

**Table S2**. Characteristics of drinking water sample before disinfectant addition.

| Characteristics | Water Sample |
|---|---|
| pH | 7.5 |
| Oxidant residual (mg $L^{-1}$ as $Cl_2$) | 0 |
| $UV_{254}$ | 0.0201 |
| $NH_3$ (mg $L^{-1}$ as N) | <0.015 |
| $NO_2^-$ (mg $L^{-1}$ as N) | <0.020 |
| $NO_3^-$ (mg $L^{-1}$ as N) | <0.1 |
| $Br^-$ (µg $L^{-1}$) | <0.1 |

**Table S3**. Free chlorine demand in each disinfection experiment. The free chlorine disinfection experiments were conducted within the same day for the same Experiment Batch; the free chlorine concentration was measured within the same day for the same Experiment Batch. The Free Chlorine Demand represents the free chlorine demand of the drinking water or coronavirus working solution after a contact time of ca. 30 s. The Free Chlorine Concentration represents the measured free chlorine concentration used in each disinfection experiment. * Negative values of free chlorine demand are resulting from the adjustment of the standard curve using a negative control of chlorine demand free phosphate buffer during each measurement.

| Experiment Batch | Water Matrix | Virus | Free Chlorine Demand (mg $L^{-1}$) | Free Chlorine Concentration (mg $L^{-1}$) |
|---|---|---|---|---|
| 1 | Drinking Water | – | 0.00 | 1.29 |
| 1 | Drinking Water | – | 0.09 | 2.51 |
| 1 | Drinking Water | MHV | -0.02* | 1.27 |
| 1 | Drinking Water | MHV | -0.01* | 2.52 |
| 2 | Phosphate Buffer | MHV | 0.04 | 1.34 |
| 3 | Phosphate Buffer | MHV | 0.06 | 2.60 |
| 4 | Drinking Water | TGEV | 0.04 | 1.27 |
| 4 | Drinking Water | TGEV | 0.13 | 2.53 |
| 5 | Phosphate Buffer | TGEV | 0.14 | 1.42 |
| 5 | Phosphate Buffer | TGEV | 0.07 | 2.38 |
| 6 | Drinking Water | HCoV-OC43 | 0.05 | 1.30 |
| 6 | Drinking Water | HCoV-OC43 | 0.25 | 2.51 |
| 7 | Phosphate Buffer | HCoV-OC43 | 0.01 | 1.27 |
| 7 | Phosphate Buffer | HCoV-OC43 | 0.17 | 2.61 |

**Table S4**. Experimental settings for free chlorine disinfection. Virus represents the target virus in a free chlorine disinfection experiment; Water Matrix represents the virus working solution matrices in free chlorine disinfection experiments; Free Chlorine Concentration represents the measured initial free chlorine concentration used in free chlorine disinfection experiments, where high and low represents the initial free chlorine concentration of 2.53±0.08 mg $L^{-1}$ and 1.31±0.06 mg $L^{-1}$ used in free chlorine disinfection experiments, representatively; Contact Time is calculated based on the Loop Length as described in the main manuscript. Three independent experiments were conducted for each condition.

| Virus | Water Matrix | Free Chlorine Concentration (mg $L^{-1}$) | Contact Time (s) |
|---|---|---|---|
| MHV | Phosphate Buffer | 1.34 | 0.55 |
| | | | 1.09 |
| | | | 2.34 |
| | | | 3.52 |
| | | 2.60 | 0.25 |
| | | | 0.55 |
| | | | 1.16 |
| | | | 1.77 |
| | Drinking Water | 1.27 | 0.55 |
| | | | 1.16 |
| | | | 2.34 |
| | | | 3.52 |
| | | 2.52 | 0.25 |
| | | | 0.55 |
| | | | 1.16 |
| | | | 1.77 |
| TGEV | Phosphate Buffer | 1.42 | 0.55 |
| | | | 1.16 |
| | | | 2.34 |
| | | | 3.52 |
| | | 2.38 | 0.25 |
| | | | 0.55 |
| | | | 1.16 |
| | | | 1.77 |
| TGEV | Drinking Water | 1.27 | 0.55 |
| | | | 1.16 |

| | | | |
|---|---|---|---|
| | | | 2.34 |
| | | | 3.52 |
| | | 2.53 | 0.25 |
| | | | 0.55 |
| | | | 1.16 |
| | | | 1.77 |
| HCoV-OC43 | Phosphate Buffer | 1.27 | 0.55 |
| | | | 1.16 |
| | | | 2.34 |
| | | | 3.52 |
| | | 2.61 | 0.25 |
| | | | 0.55 |
| | | | 1.16 |
| | | | 1.77 |
| HCoV-OC43 | Drinking Water | 1.30 | 0.55 |
| | | | 1.16 |
| | | | 2.34 |
| | | | 3.52 |
| | | 2.51 | 0.25 |
| | | | 0.55 |
| | | | 1.16 |
| | | | 1.77 |

**Table S5**. Summary of decay rate constants $k_{decay}$ of HCoV-OC43, MHV, and TGEV in drinking water and phosphate buffer. The decay rate constants $k_{decay}$ and their corresponding standard error were estimated by fitting a first-order decay model to the corresponding persistence data. The goodness of fit of the first-order decay model was assessed through examination of the coefficient of determination ($R^2$). $T_{99}$ represents the time required for a 99% reduction in viral infectivity in persistence experiments with the unit of "day".

| Water Matrix | Virus | $k_{decay}$ ($day^{-1}$) | Standard Error ($day^{-1}$) | $R^2$ | $T_{99}$ (day) |
|---|---|---|---|---|---|
| Drinking Water | MHV | 2.25 | 0.09 | 0.9938 | 2.0 |
| Phosphate Buffer | MHV | 0.28 | 0.03 | 0.8960 | 16.4 |
| Drinking Water | TGEV | 0.65 | 0.06 | 0.9400 | 7.1 |
| Phosphate Buffer | TGEV | 0.24 | 0.02 | 0.9116 | 19.2 |
| Drinking Water | HCoV-OC43 | 0.99 | 0.12 | 0.9241 | 4.7 |
| Phosphate Buffer | HCoV-OC43 | 0.51 | 0.10 | 0.8009 | 9.0 |

**Table S6**. Analysis of covariance (ANCOVA) to compare the decay rate constants of different coronaviruses in drinking water versus phosphate buffer. DW represents Drinking Water; PB represents Phosphate Buffer. For example, MHV-DW represents the decay of MHV in drinking water. p values <0.05 represent that two decay rate constants are significantly different while p values ≥0.05 represent no significant difference between two decay rate constants.

| | | MHV | | TGEV | | HCoV-OC43 | |
|---|---|---|---|---|---|---|---|
| | p | DW | PB | DW | PB | DW | PB |
| MHV | DW | | <0.0001 | <0.0001 | | <0.0001 | |
| | PB | | | | 0.5724 | | 0.0001 |
| TGEV | DW | | | | <0.0001 | <0.0001 | |
| | PB | | | | | | <0.0001 |
| HCoV-OC43 | DW | | | | | | <0.0001 |
| | PB | | | | | | |

**Table S7**. Analysis of covariance (ANCOVA) to compare the inactivation rate constants of coronaviruses by free chlorine in drinking water versus phosphate buffer, with low or high initial free chlorine concentration. Low or High represents the free chlorine concentration level. DW represents Drinking Water; PB represents Phosphate Buffer. For example, MHV-DW-Low represents the inactivation of MHV in drinking water by lower concentration of free chlorine. p values <0.05 represent that two inactivation rate constants are significantly different while p values ≥0.05 represent no significant difference between two inactivation rate constants.

| | | | MHV | | | |
|---|---|---|---|---|---|---|
| | | | DW | | PB | |
| | | p | Low | High | Low | High |
| MHV | DW | Low | | 0.9637 | 0.9522 | 0.8789 |
| | | High | | | 0.9205 | 0.8604 |
| | PB | Low | | | | 0.9194 |
| | | High | | | | |

| | | | TGEV | | | |
|---|---|---|---|---|---|---|
| | | | DW | | PB | |
| | | p | Low | High | Low | High |
| TGEV | DW | Low | | 0.9549 | 0.9380 | 0.6814 |
| | | High | | | 0.8819 | 0.7006 |
| | PB | Low | | | | 0.5970 |
| | | High | | | | |

| | | | HCoV-OC43 | | | |
|---|---|---|---|---|---|---|
| | | | DW | | PB | |
| | | p | Low | High | Low | High |
| HCoV-OC43 | DW | Low | | 0.7041 | 0.9309 | 0.6925 |
| | | High | | | 0.6099 | 0.4106 |
| | PB | Low | | | | 0.7352 |
| | | High | | | | |

**Table S8**. Summary of inactivation rate constants *kinactivate* of HCoV-OC43, MHV, and TGEV by free chlorine disinfection. The inactivation rate constants $k_{inactivate}$ and their corresponding standard error were estimated by fitting a pseudo-first-order model to the corresponding free chlorine disinfection data. The goodness of fit of the pseudo-first-order model was assessed through examination of the coefficient of determination ($R^2$).

| Virus | $k_{inactivate}$ ($mg^{-1}$ $min^{-1}$ L) | Standard Error ($mg^{-1}$ $min^{-1}$ L) | $R^2$ |
|---|---|---|---|
| MHV | 81.33 | 4.90 | 0.8284 |
| TGEV | 113.50 | 7.50 | 0.7979 |
| HCoV-OC43 | 59.42 | 4.41 | 0.7579 |

**Table S9**. Analysis of covariance (ANCOVA) to compare the inactivation rate constants of coronaviruses by free chlorine: HCoV-OC43 versus MHV versus TGEV. p values <0.05 represent that two inactivation rate constants are significantly different while p values ≥0.05 represent no significant difference between two inactivation rate constants

| p | MHV | TGEV | HCoV-OC43 |
|---|---|---|---|
| MHV | | 0.0006 | 0.0012 |
| TGEV | | | <0.0001 |
| HCoV-OC43 | | | |

**Table S10**. Akaike Information Criteria (AIC) values for the pseudo-first-order model and the biphasic model fitting the free chlorine disinfection data.

| Virus | Water | Chlorine Concentration | AIC | |
|---|---|---|---|---|
| | | | Pseudo-first-order model | Biphasic model |
| MHV | Drinking Water | low | 17.86862 | 34.48749 |
| MHV | Phosphate Buffer | low | 16.06884 | 34.98412 |
| MHV | Drinking Water | high | 20.6271 | 34.8141 |
| MHV | Phosphate Buffer | high | 13.2819 | 34.82052 |
| TGEV | Drinking Water | low | 23.59587 | 38.15368 |
| TGEV | Phosphate Buffer | low | 22.06874 | 38.63892 |
| TGEV | Drinking Water | high | 22.26501 | 38.18826 |
| TGEV | Phosphate Buffer | high | 24.4494 | 39.2188 |
| HCoV- OC43 | Drinking Water | low | 18.74666 | 32.5055 |
| HCoV- OC43 | Phosphate Buffer | low | 16.86464 | 31.66035 |
| HCoV- OC43 | Drinking Water | high | 16.5952 | 33.16959 |
| HCoV- OC43 | Phosphate Buffer | high | 17.67309 | 30.97738 |

**Table S11**. Genome composition of HCoV-OC43, MHV, and TGEV. G, C, and A are reported chlorine-reactive nucleotides in RNA genomes.[1,2] "bp" is "base pair". The complete viral genome sequences are obtained from the NIH genetic sequence database of GenBank.

| Virus | Accession Number | Genome Size (bp) | G (bp) | U (bp) | C (bp) | A (bp) | Portion of chlorine-reactive bases (G+C+A) (%) |
|---|---|---|---|---|---|---|---|
| HCoV-OC43 | AY585228.1 | 30741 | 4660 | 8502 | 6649 | 10930 | 72 |
| MHV-A59 | FJ884687.1 | 31277 | 5606 | 8094 | 7474 | 10103 | 74 |
| TGEV | AJ271965.2 | 28586 | 4847 | 8429 | 5900 | 9410 | 71 |

**Table S12**. Number of reactive amino acid residues in HCoV-OC43, MHV, and TGEV proteins, and literature values for second-order rate constants of individual amino acids, including Met (Methionine), Cys (Cysteine), His (Histidine), Trp (Tryptophan), Lys (Lysine), and Tyr (Tyrosine), reaction with HOCl at pH 7.4, 22 °C[3]. Gene Location refers to the specific position within the viral genome where the gene responsible for transcribing the corresponding viral proteins is situated. The genome sequences are obtained from UniProt database.

| Virus | Protein | Gene Location | UniProt Accession | Amino Acids (second-order reaction rate constant, $M^{-1}$ $s^{-1}$) | | | | | | Total Amino Acids |
|---|---|---|---|---|---|---|---|---|---|---|
| | | | | Met ($3.8\times10^{7}$) | Cys ($3.0\times10^{7}$) | His ($1.0\times10^{5}$) | Trp ($1.1\times10^{4}$) | Lys ($5.0\times10^{3}$) | Tyr ($4.4\times10^{1}$) | |
| HCoV-OC43 | Nucleoprotein | N | P33469 | 8 | 2 | 4 | 5 | 28 | 15 | 448 |
| HCoV-OC43 | Hemagglutinin-esterase | HE | P30215 | 6 | 14 | 7 | 4 | 13 | 36 | 424 |
| HCoV-OC43 | Envelope small membrane protein | E | Q04854 | 3 | 4 | 0 | 1 | 2 | 5 | 84 |
| HCoV-OC43 | Membrane protein | M | Q01455 | 10 | 3 | 3 | 7 | 11 | 14 | 230 |
| HCoV-OC43 | Spike glycoprotein | S | P36334 | 21 | 55 | 13 | 14 | 59 | 78 | 1353 |
| TGEV | Nucleoprotein | N | P04134 | 3 | 2 | 5 | 7 | 35 | 10 | 382 |
| TGEV | Envelope small membrane protein | E | P09048 | 4 | 3 | 1 | 1 | 3 | 3 | 82 |
| TGEV | Membrane protein | M | P04135 | 9 | 8 | 2 | 10 | 13 | 15 | 262 |
| TGEV | Spike glycoprotein | S | P07946 | 14 | 49 | 22 | 23 | 45 | 67 | 1447 |
| MHV-A59 | Nucleoprotein | N | P03416 | 4 | 2 | 4 | 5 | 32 | 11 | 454 |

| | | | | | | | | | | |
|---|---|---|---|---|---|---|---|---|---|---|
| MHV-A59 | Envelope small membrane protein | E | P0C2R0 | 3 | 4 | 0 | 1 | 3 | 6 | 83 |
| MHV-A59 | Membrane protein | M | P03415 | 8 | 4 | 3 | 7 | 10 | 13 | 228 |
| MHV-A59 | Spike glycoprotein | S | P11224 | 14 | 53 | 18 | 14 | 52 | 68 | 1324 |

**Table S13**. Correlation matrix between virus inactivation rate constants by free chlorine disinfection and viral protein composition. S, N, M, and E represent spike protein, nucleoprotein, membrane protein, and envelope small membrane protein, representatively, in HCoV-OC43, MHV, and TGEV. Met, Cys, His, Trp, Lys, and Tyr represent the absolute counts of corresponding amino acids in spike protein, nucleoprotein, membrane protein or envelope small membrane protein. *k* represents the coronavirus inactivation rate constants by free chlorine disinfection estimated by fitting a pseudo-first-order model to the corresponding free chlorine disinfection data. The correlation coefficients are shown as digital numbers and the significance level are shown as stars. "***", "**", "*", ".", and " " represent p-values of 0, 0.001, 0.01, 0.05, 0.1, and 1 respectively. NA represents not applicable.

| correlation coefficients | Met | Cys | His | Trp | Lys | Tyr |
|---|---|---|---|---|---|---|
| | S Protein | | | | | |
| k | -0.74*** | -0.86*** | 0.87*** | 0.77*** | -0.87*** | -0.78*** |
| | N Protein | | | | | |
| k | -0.82*** | NA | 0.77*** | 0.77*** | 0.87*** | -0.82*** |
| | M Protein | | | | | |
| k | -0.41* | 0.83*** | -0.77*** | 0.77*** | 0.59*** | 0.46** |
| | E Protein | | | | | |
| k | 0.77*** | -0.77*** | 0.77*** | NA | 0.74*** | -0.59*** |
